# Support for the dominance theory in *Drosophila* transcriptomes

**DOI:** 10.1101/321455

**Authors:** Ana Llopart, Evgeny Brud, Nikale Pettie, Josep M. Comeron

## Abstract

Interactions among divergent elements of transcriptional networks from different species can lead to misexpression in hybrids through regulatory incompatibilities, some with the potential to generate sterility. Genes with male-biased expression tend to be overrepresented among genes misexpressed in hybrid males. While the possible contribution of faster-male evolution to this misexpression has been explored, the role of the hemizygous *X* chromosome (i.e., the dominance theory for transcriptomes) remains yet to be determined. Here we study genome-wide patterns of gene expression in females and males of *Drosophila yakuba* and *D. santomea* and their hybrids. We used attached-X stocks to specifically test the dominance theory, and we uncovered a significant contribution of recessive alleles on the *X* chromosome to hybrid misexpression. Our analysis of gene expression patterns suggests that there is a contribution of weakly deleterious regulatory mutations to gene expression divergence in the sex towards which the expression is biased. In the opposite sex (e.g., genes with female-biased expression analyzed in male transcriptomes), we detect stronger selective constraints on gene expression divergence. Although genes with high degree of male-biased expression show a clear signal of faster-X evolution for gene expression divergence, we also detected slower-X evolution of gene expression in other gene classes (e.g. female-biased genes) that is mediated by significant decreases of *cis*- and *trans*-regulatory divergence. The distinct behavior of X-linked genes with high degree of male-biased expression is consistent with these genes experiencing a higher incidence of positively selected regulatory mutations than their autosomal counterparts. We propose that both dominance theory and faster-X evolution of gene expression may be major contributors to hybrid misexpression and possibly the large X-effect in these species.

## INTRODUCTION

Population genetics models predict that, under certain conditions, X-linked loci are expected to show accelerated divergence relative to autosomal loci (i.e., faster-X evolution) (Charlesworth *et al.* 1987; Orr and Betancourt 2001; Betancourt *et al.* 2002; Vicoso and Charlesworth 2006; Vicoso and Charlesworth 2009a; Mank *et al.* 2010b). Faster-X evolution can, in principle, result from the contributions to the rates of evolution of either beneficial or weakly deleterious mutations. The visibility to selection of new X-linked recessive beneficial mutations in hemizygous males significantly increases their chances of spreading and contributing to the rate of evolution. Weakly deleterious mutations that are partially dominant can also produce faster-X because the efficacy of natural selection tends to be slightly reduced on the *X* chromosome. This is because the effective population size of the *X* chromosome (*N_eX_*) is often reduced relative to that of autosomes (*N_eA_*) due to its hemizygosity in males. Theoretical models, however, show that the chances of observing faster-X evolution are very sensitive to the ratio *N_eX_* /*N_eA_* (Vicoso and Charlesworth 2009a; Mank *et al.* 2010b) and any parameter that impacts this ratio, including demographic changes and differences in recombination rates between the *X* chromosome and autosomes among others (Langley *et al.* 1988; Connallon 2007; Pool and Nielsen 2007; Vicoso and Charlesworth 2009b; Charlesworth 2012; Comeron *et al.* 2012; Avila *et al.* 2015). For extreme values of *N_eX_* /*N_eA_*, faster-X evolution can even occur independently of the coefficient of dominance (Vicoso and Charlesworth 2009a; Mank *et al.* 2010b).

Today there is ample evidence supporting faster-X evolution of protein-coding sequences in a variety of organisms, including *XY* and *ZW* systems (Thornton and Long 2002; Torgerson and Singh 2003; Counterman *et al.* 2004; Khaitovich *et al.* 2005; Lu and Wu 2005; Nielsen *et al.* 2005; Thornton and Long 2005; Musters *et al.* 2006; Torgerson and Singh 2006; Baines and Harr 2007; Begun *et al.* 2007; Mank *et al.* 2007; Baines *et al.* 2008; Singh *et al.* 2008; Ellegren 2009; Mank *et al.* 2010a; Grath and Parsch 2012; Meisel and Connallon 2013; Vicoso *et al.* 2013; Avila *et al.* 2014; Garrigan *et al.* 2014; Kousathanas *et al.* 2014; Sackton *et al.* 2014; Veeramah *et al.* 2014; Avila *et al.* 2015; Coolon *et al.* 2015; Llopart 2015; Larson *et al.* 2016). Faster-X evolution has also been reported for gene expression divergence (Khaitovich *et al.* 2005; Brawand *et al.* 2011; Kayserili *et al.* 2012; Llopart 2012; Meisel *et al.* 2012; Coolon *et al.* 2015; Dean *et al.* 2015) and intergenic DNA sequences (Hu *et al.* 2013; Coolon *et al.* 2015; Llopart 2018), likely reflecting the effects of natural selection on the evolution of regulatory elements. In species where males are the heterogametic sex, faster-X is expected to be strongest in genes with male-specific fitness effects because these genes are always hemizygous when they are X-linked. A similar rationale applies to genes with female-specific fitness effects in *ZW* species (Betancourt *et al.* 2002; Vicoso and Charlesworth 2009a; Mank *et al.* 2010b).

Regardless of the evolutionary forces driving faster-*X* (*Z*) evolution (positive selection or relaxed negative selection), its implications for the genetic basis of hybrid dysfunction are clear: the *X* chromosome will disproportionally contribute to between-species Dobzhansky-Muller incompatibilities (Dobzhansky 1937; Muller 1942; Coyne and Orr 1989). Even a slightly elevated substitution rate on the *X* chromosome will be amplified in hybrids, as the number of incompatibilities increases exponentially with divergence time (Orr 1995; Turelli and Orr 2000; Matute *et al.* 2010; Moyle and Nakazato 2010; Guerrero *et al.* 2017). Mapping studies in a variety of species including mice, *Lepidoptera*, and *Drosophila* have shown that the factors having the largest effects on hybrid dysfunction, particularly hybrid male sterility, are X-linked (Coyne and Orr 1989; Coyne *et al.* 1991; Coyne 1992; Coyne and Orr 2004). Orr’s (1987) seminal study of the genetic basis of hybrid sterility in *D. pseudoobscura-D. persimilis* constitutes a classic example of this pattern known as the large X-effect. The large X-effect, together with Haldane’s rule (i.e., the preferential inviability or sterility of the heterogametic over the homogametic sex; Haldane 1922), underline the importance of sex chromosomes in speciation (Coyne and Orr 1989; Masly and Presgraves 2007; Presgraves 2008; Moyle *et al.* 2010).

The large X-effect can also be explained by the dominance theory (Turelli and Orr 1995; Orr and Turelli 1996; Turelli and Orr 2000), which posits that alleles causing hybrid dysfunction tend to act recessively in hybrids (Muller 1940;Muller 1942). As a result, hemizygous *X* chromosomes will express all incompatibilities (dominant and recessive) while heterozygous autosomes will only express the fairly dominant ones. In the context of gene regulation, the dominance theory, if confirmed, opens the possibility that a large fraction of the the abundant misexpression observed in hybrid males, but not in hybrid females, may be due to regulatory incompatibilities involving X-linked recessive mutations. Disruption of gene expression has been extensively documented in *Drosophila* hybrids and it is often the case that rapidly evolving genes enriched in male functions tend to be overrepresented among misexpressed genes in sterile hybrid males (Reiland and Noor 2002; Michalak and Noor 2003; Michalak and Noor 2004; Ranz *et al.* 2004; Landry *et al.* 2005; Noor 2005; Haerty and Singh 2006; Artieri *et al.* 2007; Barbash and Lorigan 2007; Moehring *et al.* 2007; Catron and Noor 2008; Sundararajan and Civetta 2011; Llopart 2012; Maheshwari and Barbash 2012; Satyaki *et al.* 2014; Wei *et al.* 2014; Gomes and Civetta 2015). While the potential link between faster-male theory (Wu and Davis 1993; Wu *et al.* 1996) and hybrid misexpression has been explored (Noor 2005; Haerty and Singh 2006; Ortiz-Barrientos *et al.* 2007; Civetta 2016), the dominance theory for hybrid transcriptomes remains untested. Here we examine genome-wide patterns of transcript abundance in females and males of *Drosophila yakuba, D. santomea* and their interspecific hybrids using RNAseq technology for genes with variable degrees of sex-biased expression. We took advantage of attached-X stocks developed in both species to test the dominance theory for transcriptomes.

## MATERIALS AND METHODS

### Fly stocks

We used the *D. yakuba* Taï18E2 and the *D. santomea* STO.4 stocks to obtain an overall view of gene expression variation between species and in F_1_ hybrid females and males. The *D. yakuba* Taï18E2 was part of the 12 *Drosophila* Genomes project, originated in the Taï rainforest (Taï National Park, Ivory Coast), and was inbred for at least 10 generations of single-pair brother/sister mating before whole-genome sequencing (Clark *et al.* 2007). The STO.4 stock of *D. santomea* was established from a single fertilized female collected in the Obo Natural Reserve on São Tomé (Lachaise *et al.* 2000), and was inbred for 10 generations. To generate ‘unbalanced’ F_1_ hybrid females with two *D. yakuba X* chromosomes, *D. yakuba* females of the attached-X stock [C(1)RM *y, w^or^* females, + males] were crossed to *D. santomea* males of the attached-X stock [C(1)RM *g* females, + males] (Coyne *et al.* 2004); F_1_ hybrid females with two *D. santomea X* chromosomes were also generated from the reciprocal cross. All flies were raised on standard cornmeal-yeast-agar medium at 24°C with a 12 h dark-light cycle.

### RNA isolation, library preparation, and sequencing

Females and males were collected under CO_2_ anesthesia 0-5 h post eclosion, aged for 19 h at 24°C, snap-frozen in liquid nitrogen, and stored at –80°C. For each genotype analyzed, total RNA was isolated from a pool of 15 females or 30 males using TRIzol Reagent and following the manufacturer’s recommendations (Invitrogen). Up to 15 μg of nucleic acid were treated with RNase-free DNase I (∼7 Kunitz units) in solution (‘RNase-free DNase set’; Qiagen) and cleaned- up using the RNeasy Mini Kit (Qiagen). We used ∼1 μg of total RNA to prepare libraries suitable for high throughput sequencing following the Illumina protocol (‘mRNA Sample Preparation Guide’; Cat. # RS-930-1001, Part # 1004898 Rev. D; Illumina). Briefly, poly-A containing RNA molecules were purified using poly-T oligo-attached magnetic beads. Following this purification, the mRNA was fragmented into small pieces using divalent cations under elevated temperature. The first-strand cDNA was copied using reverse transcriptase and random hexamers while the second cDNA strand was synthesized using DNA polymerase I and RNAse H. The fragmented cDNA went through the end repair process, addition of a single ‘A’ base, and ligation to custom-designed adapters. These include unique 7-nucleotide tags that allowed multiplexing of several samples prior to sequencing. The sequences and a detailed description of these custom-designed adapters can be found elsewhere (Comeron *et al.* 2012).

We run the ligation products on a 2% agarose gel, selected for ∼250bp-size, and recovered cDNA templates using the QIAquick Gel Extraction Kit (Qiagen). These cDNA templates were enriched with PCR to create the final cDNA libraries, which were validated using a 2100 Bioanalyzer (Agilent Technologies). For each genotype analyzed, we obtained two technical replicates using the same total RNA. Cluster generation and sequencing were carried out on HiSeq 2000/2500 instruments at the Iowa Institute of Human Genetics (IIHG; University of Iowa) or at the Iowa State DNA facility (Iowa State University) using a combination of multiple lanes per sample (single-end 100 cycles/read or 125 cycles/read). Sequence reads were sorted out according to individual tags, 3’ trimmed (-t 12 -l 30), and filtered based on quality (-q 12 and -p 75) using the FASTX-toolkit (0.0.14) (http://hannonlab.cshl.edu/fastx_toolkit/commandline.html). The average number of trimmed reads that passed quality filtering across samples was 51.5 and 31.3 millions for replicates 1 and 2, respectively. All analyses were carried out combining sequence reads from both technical replicates. All sequence reads have been archived at NCBI SRA BioProject PRJNA470895.

### Gene expression analyses

To examine genome-wide transcript abundance, we first generated the *D. yakuba* reference gene set (14,687 genes) based on the genome sequence and annotation of coding sequences (CDS) available in FlyBase (dyak_r1.3_FB2011_08; http://flybase.org/) (Gramates *et al.* 2017). Genes on Arm U and random sequence scaffolds were not included. To allow reads covering 5’ and 3’ UTRs, we expanded the CDS definitions by 75 base pairs upstream and downstream of starting and stop codons, respectively. Exon and UTR sequences were extracted using tools available on Galaxy (https://usegalaxy.org/) (Afgan *et al.* 2016) and assembled with custom scripts.

Trimmed and quality filtered reads were mapped to the *D. yakuba* reference gene set with Bowtie 2 (version 2.1.0) using default parameters(Langmead and Salzberg 2012). Bam files were sorted and indexed with SAMtools [version 0.1.18 (r982:295)] (Li *et al.* 2009), and the number of reads aligned to each of the genes was calculated using RSEM (version 1.3.0) (Li and Dewey 2011). We only considered for further analyses genes with at least 10 mapped reads and > 1 fpkm (Fragments Per Kilobase of transcript per Million mapped reads) in both species and in at least one sex.

To classify genes into different sex-biased expression categories that were consistent across species, we took advantage of a DESeq2 (version 1.16.1) multifactor design with two variables (∼species + sex) and used the effect size estimate (Log2FoldChange) as a measure of the degree of sex-biased expression (Love *et al.* 2014). Genes showing a fold-change difference in expression between the two sexes smaller than 1.1 were classified as nonsex-biased. Genes showing statistically significant differences in expression between females and males and < 1 fpkm in one sex were considered to have sex-specific expression. Specifically, we detected 156 genes expressed in females but not in males and 1,660 genes showing the opposite pattern. Quantitative estimates of differences in gene expression between species, between parental species and hybrids, and between reciprocal interspecific hybrids, were also based on size effects from DESeq2. DESeq2 implements the Benjamini-Hochberg probability adjustment (padj) to correct for multiple testing (Benjamini and Hochberg 1995). Genes were considered differentially expressed if padj < 0.05 (FDR = 0.05). We excluded genes showing switches across species between sex-biased expression categories (e.g., genes with male-biased expression in *D. yakuba* but female-biased expression in *D. santomea*). We analyzed gene expression for a total of 11,959 genes.

### Analyses of *cis*- and *trans*-regulatory variation

The detection of *cis*- and/or *trans*-regulatory variation is usually based on comparing the ratio of allelic expression in hybrids (i.e., allelic ratio) with the ratio of expression between species (i.e., species ratio) (Wittkopp *et al.* 2004; McManus *et al.* 2010; Coolon *et al.* 2014). With mRNAseq data, this requires the identification of parental stock-specific reads, which was based on whole-genome consensus sequences for the *D. yakuba* Taï18E2, the *D. santomea* STO.4, the *D. yakuba* attached-X, and the *D. santomea* attached-X stocks. The initial mapping steps for generating these new genome sequences were the same as those used for the analysis of transcript abundance (see above) except that, in this case, we used the *D. yakuba* reference genome sequence to align *D. yakuba* reads and a draft of the *D. santomea* genome sequence (Llopart 2015)to align *D. santomea* reads. SAMtools was used to (1) sort/index BAM files, (2) filter sites with mapping quality < 40, and (3) generate mpileup files with a maximum depth of 100,000 reads and a minimum base quality of 30 (Li *et al.* 2009). To further filter sites covered by fewer than 5 reads and to obtain fastq files of consensus sequences, we used the utility BCFtools (1.3.1) (Li *et al.* 2009). Fastq files for each of the four consensus sequences were converted into fasta format and heterozygous sites were randomly phased using Heng Li’s seqtk (https://github.com/lh3/seqtk). To minimize mapping biases across comparisons, regions containing ambiguous bases or missing base calls in at least one of the consensus sequences were also masked in all the other consensus sequences.

Based on comparisons between allelic and species ratios, we classified genes into six different regulatory evolution classes (*cis* only, *trans* only, *cis* + *trans, cis* × *trans*, compensatory, conserved) following McManus *et al*. (2010). For each type of hybrid analyzed, we identified stock-specific reads as those mapping with 0 mismatches to only one of the parental consensus sequences. The relative numbers of the two types of stock-specific reads in hybrids provide an estimate of the allelic ratio of gene expression, which was calculated as log_2_(read count allele 1/ read count allele 2). This ratio constitutes a quantitative measure of *cis*-regulatory divergence for every gene analyzed. To obtain an estimate of the species ratio of expression, we first constructed an *in silico* parental mixture by combining equal numbers of stock-specific reads from each of the two parental stocks used to generate the different hybrid types, as in Coolon *et al*. (2014). The species ratio was then calculated as log_2_(read count genotype 1/ read count genotype 2) (Coolon *et al.* 2014; Coolon *et al.* 2015). The difference between species and allelic ratios [log_2_(read count genotype 1/ read count genotype 2) - log_2_(read count allele 1/read count allele 2)] provides a quantitative measure of *trans*-regulatory divergence for each gene analyzed. Binomial exact tests were used to detect differences between read counts of allele 1 and allele 2, and also between read counts of genotype 1 and genotype 2. A χ^2^ test was applied to detect differences between allelic and species ratios. Sequential Bonferroni correction for multiple testing was used (FDR = 0.05). Only genes with at least 20 stock-specific mapped reads for the two types of alleles and genotypes were included in the analyses.

## RESULTS

To understand the role of the *X* chromosome in hybrid gene expression, we examined transcript abundance in 1-day old virgin females and males of *Drosophila yakuba, D. santomea* and their interspecific hybrids. Our experimental design includes not only flies with normal karyotypes but also females who carry attached-X chromosomes (Coyne *et al.* 2004) (**Figure** 1). Attached-X hybrid females have two *X* chromosomes from the same species and a haploid set of autosomes from each parental species. They are as ‘unbalanced’ as hybrid males for the *X*/Autosome ratio and constitute an ideal genetic tool to study the effects of X-linked recessive factors. We obtained profiles of transcript abundance for 10,128 and 11,663 genes expressed in adult females and males, respectively. To investigate the effects of sexually dimorphic gene expression, we classified all genes into seven distinct categories based on degree of sex-biased expression (**Figure** 2). Importantly, this gradual classification allows for comparisons across studies regardless of statistical power. In addition and to capture extreme sex-biased effects, we also identified genes expressed in females but not in males (female-specific genes, FSGs) and genes showing the opposite pattern (male-specific genes, MSGs).

**Figure 1.**
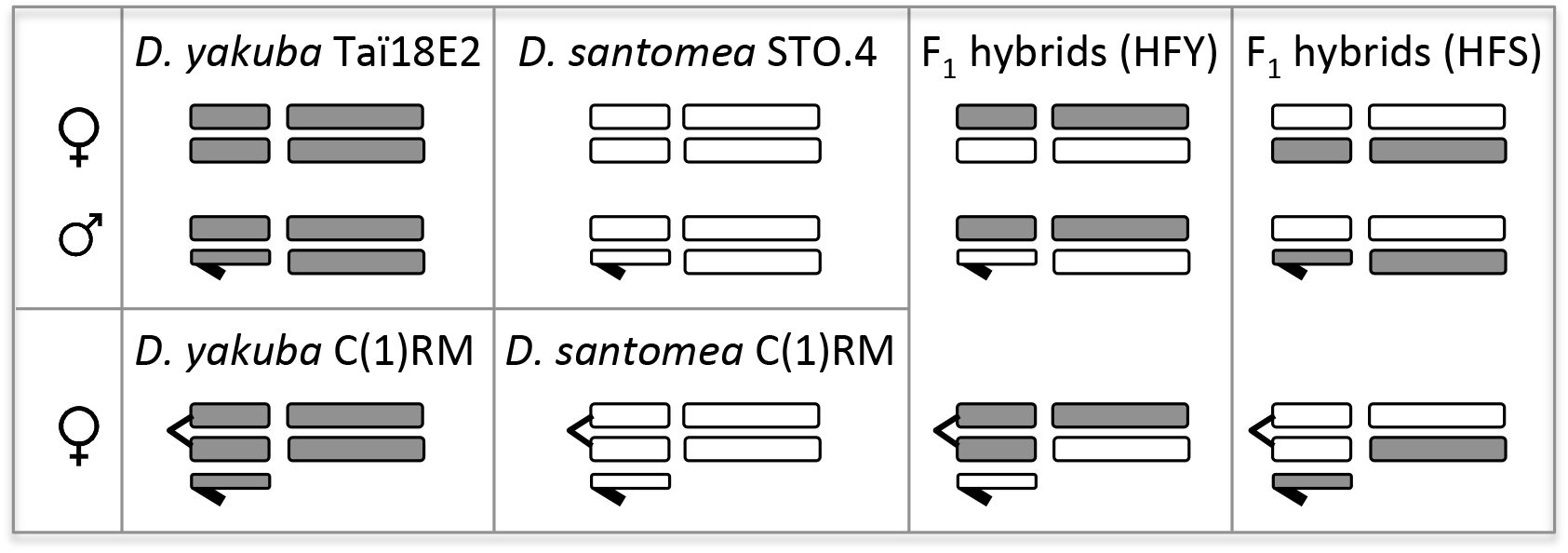
Experimental design using attached-X females and karyotypes analyzed. Horizontal short bars represent sex chromosomes (the *Y* chromosome is shown with a hook) while long bars represent autosomes. Grey, *D. yakuba* chromosomes; white, *D. santomea* chromosomes. *HFY*, hybrids of first generation from the cross between *D. yakuba* (Taï18E2 or attached-X stocks) females and *D. santomea* (STO.4 or attached-X stocks) males; *HFS*, hybrids of first generation from the cross between *D. santomea* (STO.4 or attached-X stocks) females and *D. yakuba* (Taï18E2 or attached-X stocks) males.

**Figure 2.**
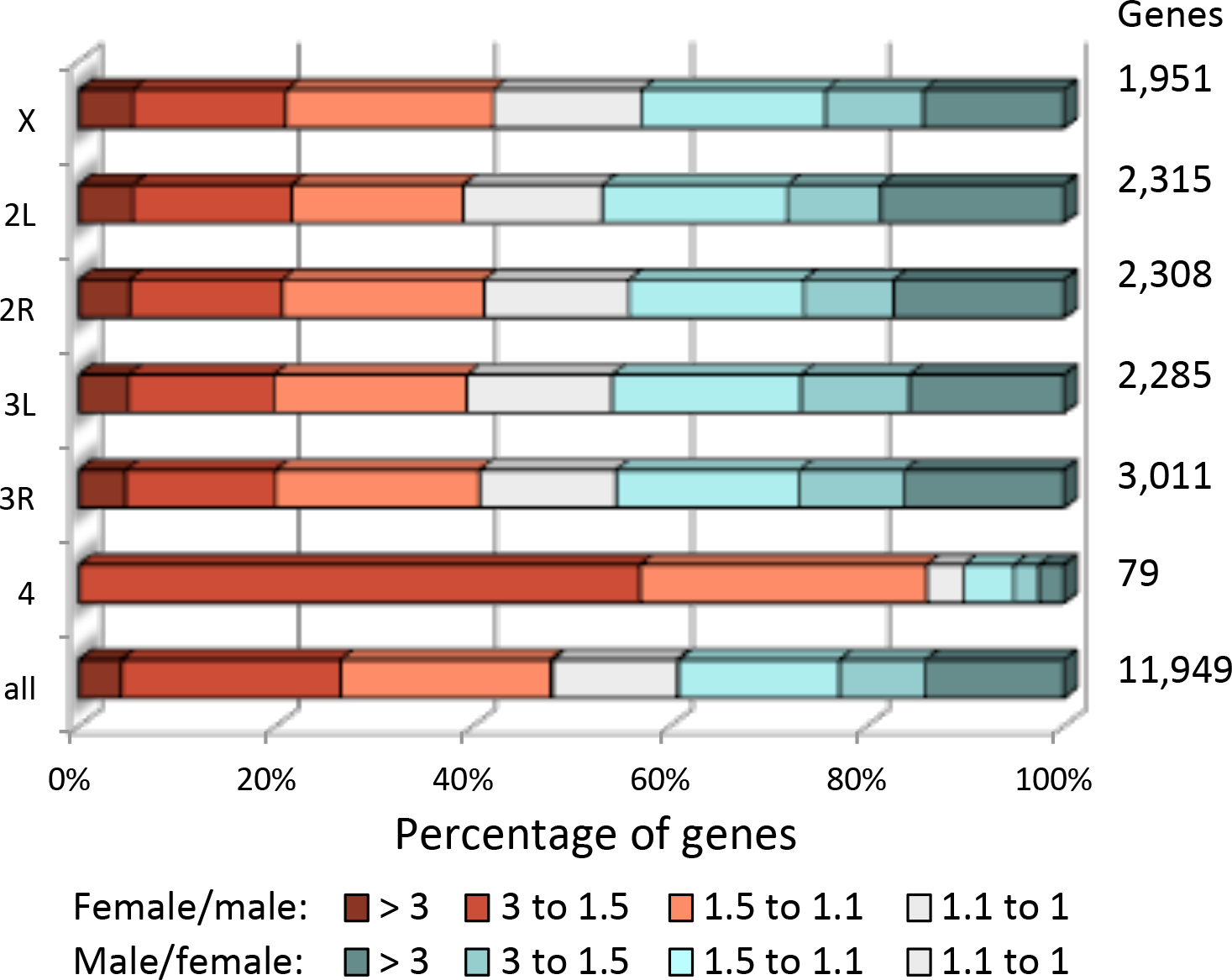
Distribution of genes with sexually dimorphic gene expression across the genome. The number of genes in each class is shown on the right column.

To investigate the relationship between sexually dimorphic expression and chromosome location in the *D. yakuba*-*D. santomea* system, we examined the genome-wide distribution of genes with variable degrees of sex-biased expression (**Figure** 2). The autosomal dot (fourth) chromosome reveals the most striking pattern, with 86% (68/79) of its genes being female-biased in expression. This represents a significant enrichment relative to the other autosomes [68/79 *vs*. 3,978/9,919; Fisher’s Exact Test (FET), *P* = 5.08×10^-17^], which we interpret as evidence of ancient feminization of the dot chromosome (Vicoso and Bachtrog 2013; Vicoso and Bachtrog 2015). The *X* chromosome shows a significant overrepresentation of FSGs relative to autosomes (40/1,951 *vs*. 116/9,919, FET *P* = 0.003) and a trend towards underrepresentation of MSGs (246/1,951 *vs*. 1,413/9,919, FET *P* = 0.058).

### Gene expression divergence is a sexually dimorphic trait

To understand the molecular causes underlying hybrid misexpression, we first examined gene expression divergence using a quantitative measure of the difference in transcript abundance between species [i.e., log_2_(*D. yakuba*-to-*D. santomea* ratio of expression)]. These analyses uncovered three main patterns. First, gene expression divergence is positively correlated with degree of sex-biased expression [Spearman’s ρ = 0.10 (95% confidence intervals: 0.07–0.13) (*P* = 1.04×10^-11^) for genes with increasingly female-biased expression in females and ρ = 0.30 (95% CI: 0.27–0.32) (*P* = 3.47×10^-111^) for genes with increasingly male-biased expression in males] (**Figure** 3). Second, female transcriptomes show smaller overall gene expression divergence relative to male transcriptomes (0.28 *vs*. 0.30; Mann-Whitney Test (MWT) *P* = 6.75×10^-12^], which is consistent with faster-male evolution (Wu and Davis 1993; Wu *et al.* 1996) for gene expression (Meiklejohn *et al.* 2003; Parisi *et al.* 2003; Ranz *et al.* 2003; Haerty *et al.* 2007). Gene expression also evolves more rapidly in MSGs than in FSGs (0.58 vs. 0.41; MWT *P* = 0.007). Third, the comparative analysis of the *same* subset of genes in the female and male transcriptomes reveals that genes with sex-biased expression show high gene expression divergence in the sex toward which the expression is biased while transcript abundance tends to be conserved between species in the opposite sex; the trend is consistently observed across genes with different degrees of either female- or male-biased expression (**Figure** 3). As a result, some gene categories can show greater gene expression divergence when expressed in females than when expressed in males. This suggests that, in addition to the pervasive trend of faster-male evolution, there is a signal for rapid evolution in the female transcriptome as well. We suggest that gene expression divergence for genes with sex-biased expression behaves as a sexually dimorphic trait.

**Figure 3.**
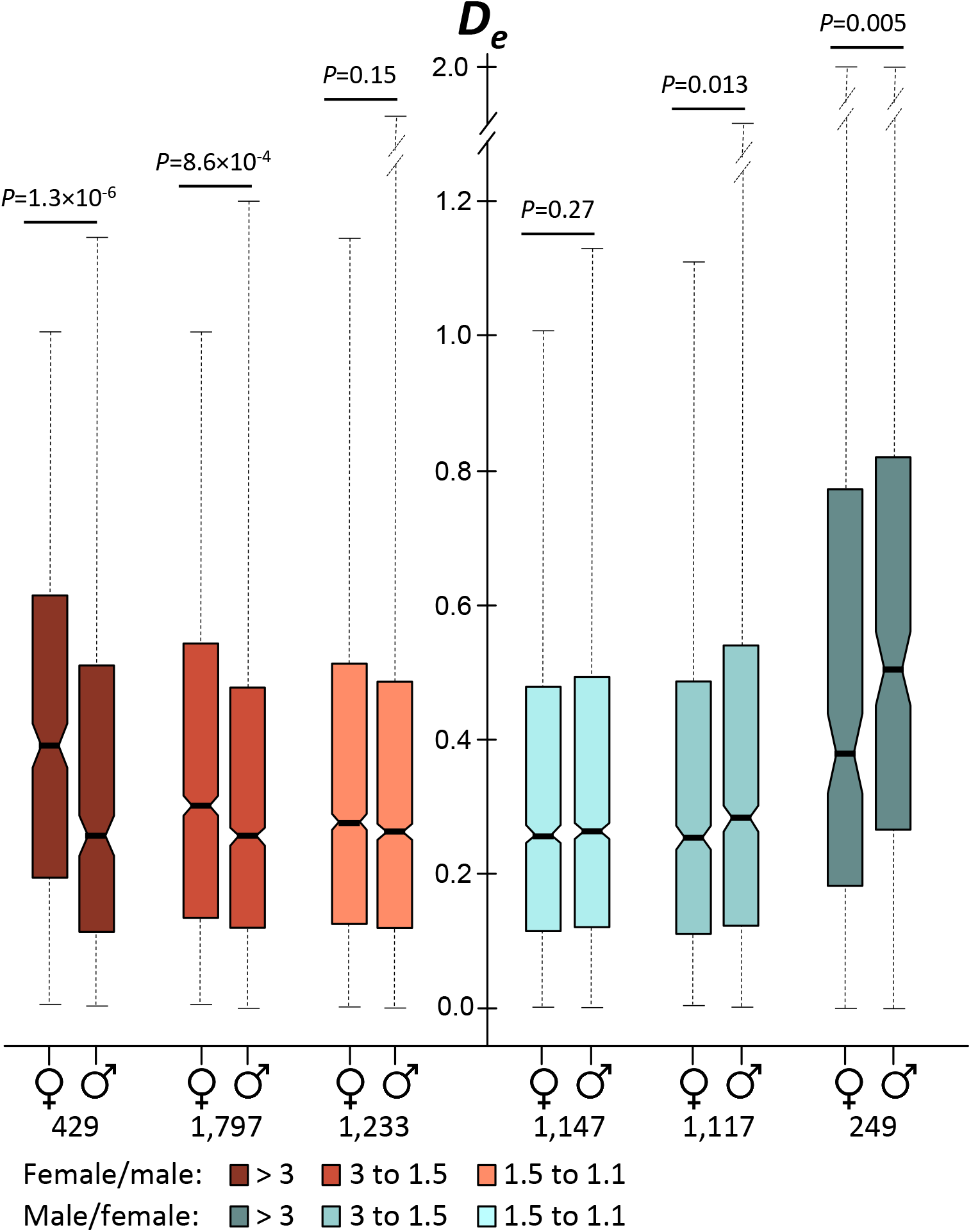
Gene expression divergence (*D_e_*) as a sexually dimorphic trait in genes with sex-biased expression. The horizontal line inside each box indicates the median. The length of the box and the whiskers represent 50% and 90% confidence intervals, respectively. The numbers of genes analyzed in each sex-biased category are shown under female and male symbols. Probabilities are based on Mann-Whitney tests.

### Weakly deleterious regulatory mutations contribute to gene expression divergence

To gain insight into the evolutionary forces overriding gene expression divergence, we analyzed transcript abundance on the dot chromosome. In *Drosophila*, the dot chromosome is a unique autosome because it does not experience meiotic recombination. Due to linked selection, the effective population size and the effectiveness of natural selection on the dot chromosome are severely reduced (Hill and Robertson 1966; Berry *et al.* 1991; Betancourt and Presgraves 2002; Comeron and Kreitman 2002; Jensen *et al.* 2002; Wang *et al.* 2002; Wang *et al.* 2004; Haddrill *et al.* 2007; Comeron *et al.* 2008; Betancourt *et al.* 2009; Arguello *et al.* 2010). Because of the highly biased gene composition of the dot chromosome (**Figure** 2), we focused our analyses in genes with female-biased expression.

If weakly deleterious mutations are major contributors to gene expression divergence, we anticipate that gene expression on the dot chromosome will evolve faster than on other autosomes. This prediction is precisely observed but only in the female transcriptome: gene expression divergence for genes with female-biased expression on the dot chromosome is significantly higher than that on other autosomes (0.62 *vs*. 0.29; MWT *P =* 9.65×10^-13^). In male transcriptomes, by contrast, there is no evidence of increased gene expression divergence for female-biased genes on the dot chromosome (0.21 *vs*. 0.27; MWT *P =* 0.06), which indicates minimal effects of differences in effective population size and thus suggest selectively strong functional constraints. The presence of only eight genes with male-biased expression on the dot chromosome precludes us from formally testing whether genes with male-biased expression, when expressed in males, show any evidence of a contribution of weakly deleterious mutations to gene expression divergence. The comparison of female and male transcriptomes, therefore, suggests that gene expression divergence is not inherently high in genes with sex-biased expression, and that there is a major contribution of weakly deleterious regulatory mutations but only in the sex toward which expression is biased.

### Faster-X evolution for gene expression divergence in highly male-biased genes

We also examined gene expression divergence in the context of the comparison between X-linked and autosomal genes. The female transcriptome shows an overall signal in support of slower-X evolution of gene expression (0.25 *vs*. 0.28, MWT *P* = 4.81×10^-5^), although for some gene categories the signal is weak or not significant (**Figure** 4A). The male transcriptome also shows slower-X evolution for gene expression divergence when all genes are considered as a single class (0.28 *vs*. 0.31, MWT *P* = 0.002). Despite this overall trend, we found significant evidence for faster-X evolution of gene expression in genes with high degree of male-biased expression (**Figure** 4B). Faster-X is particularly pronounced in MSGs (0.81 *vs*. 0.54, MWT *P* = 4.13×10^-13^). X-linked MSGs also show the greatest gene expression divergence of all gene classes in the male transcriptome, including the fastest evolving genes on the dot chromosome (0.81 *vs*. 0.68, MWT *P* = 0.017).

**Figure 4.**
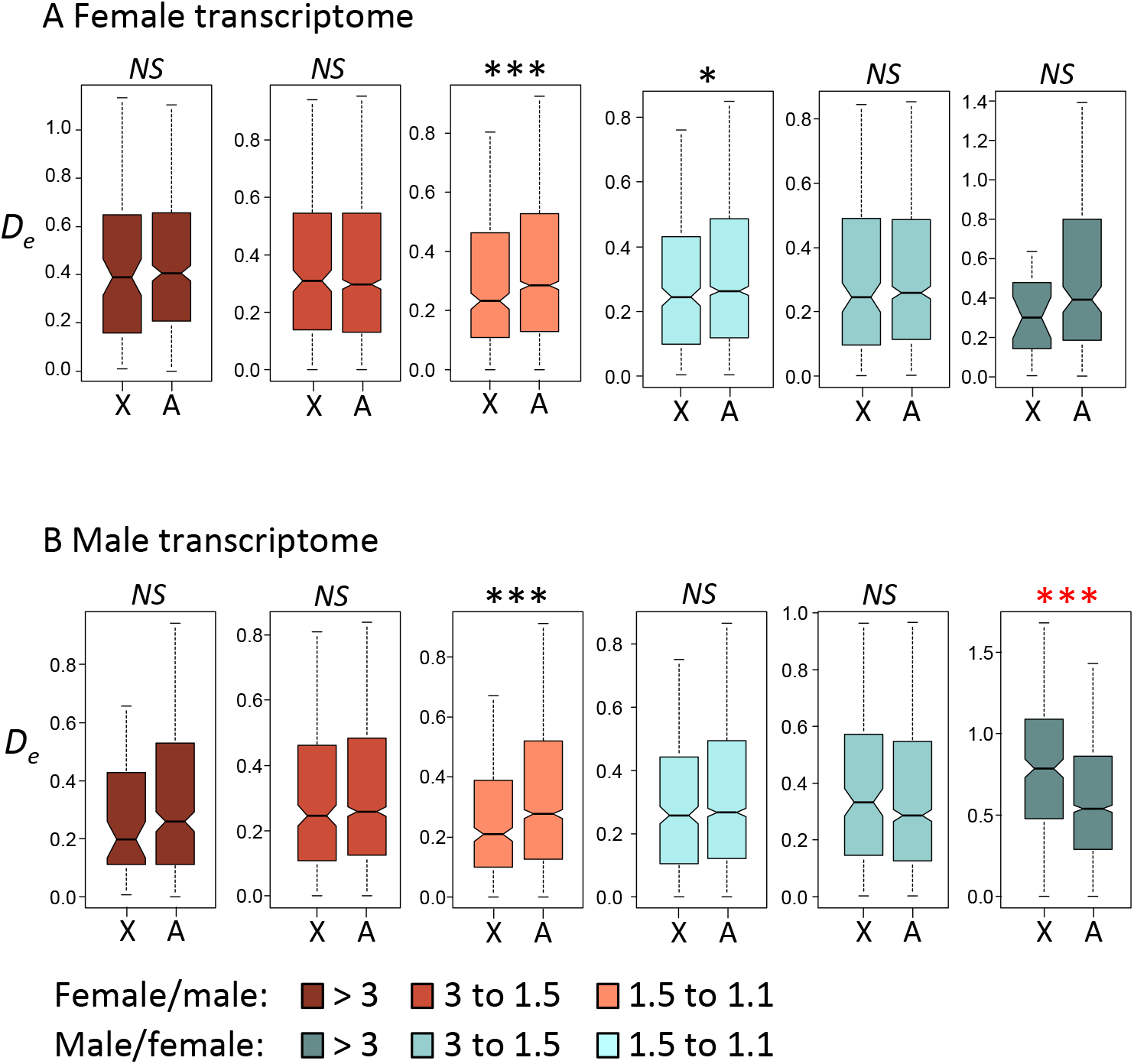
Tests of faster-X evolution for gene expression divergence in the female transcriptome (A) and the male transcriptome (B). Black and red asterisk indicate slower-X and faster-X, respectively. Probabilities are based on Mann-Whitney tests where *NS* indicates not significant, \**P* < 0.05, \*\**P* < 0.001, and \*\*\**P* < 1×10^-4^. (See Fig. 3 legend for boxplot explanation.)

To gain insight into the genetic basis (*cis*- or *trans*-regulatory) of gene expression divergence, we examined stock-specific expression following Coolon *et al*. (2015) (Wittkopp *et al.* 2004; McManus *et al.* 2010; Coolon *et al.* 2014) (see also ‘Materials and Methods’). We found strong positive correlations between gene expression divergence and the magnitude of both *cis*- and *trans*-regulatory divergence (Table 1). These correlations imply that any increase in gene expression divergence is likely to be the result of contributions from *cis*- and *trans*-regulatory variation. For example, the overall trend of slower-X evolution for gene expression divergence in females is accompanied by significantly greater *cis*- (0.26 *vs*. 0.28; MWT *z* = 2.82, *P* = 0.005) and *trans*-regulatory (0.19 *vs*. 0.23; MWT z = 5.04, *P* = 4.8×10^-7^) divergence for autosomal than for X-linked genes. Likewise autosomal genes in the male transcriptome show increased *cis*- (0.30 *vs*. 0.28, MWT *z* = 3.22, *P* = 0.0012) and *trans*-regulatory (0.24 *vs*. 0.23, MWT *z* = 3.36, *P* = 0.0008) divergence relative to the same genes when analyzed in the female transcriptome, as expected from faster-male evolution. As a result, when we examined the different categories of genes with sexually dimorphic expression, we detected that the fraction of genes with significant *cis*-regulatory divergence increases with gene expression divergence; the same was observed for *trans*-regulatory divergence.

**Table 1.**
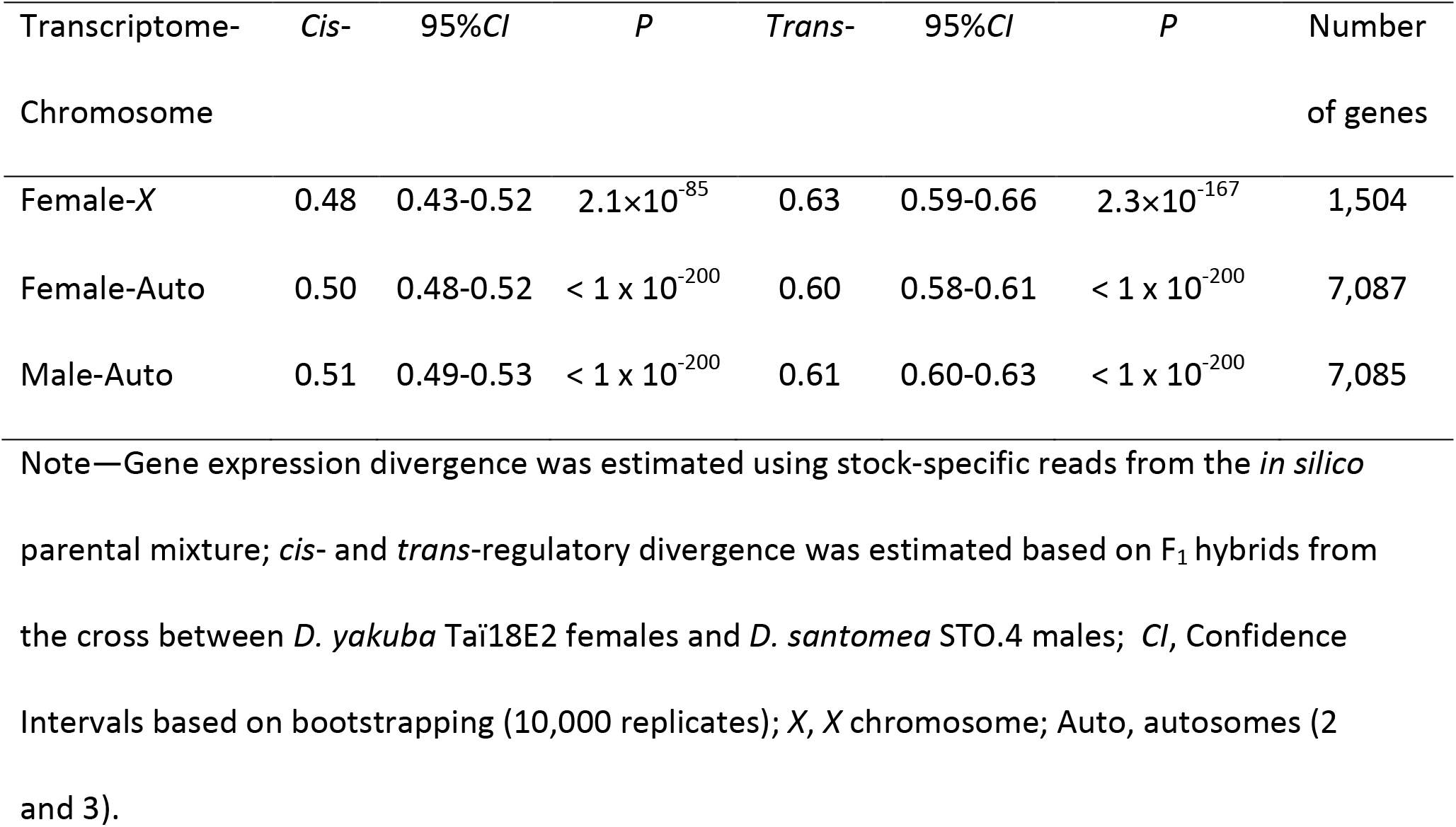
Spearman’s correlations (ρ) between gene expression divergence and the magnitude of *cis*- and *trans*-regulatory divergence

### Recessive regulatory incompatibilities involving the *X* chromosome contribute to hybrid misexpression

To test the dominance theory (Muller 1940; Muller 1942) for hybrid transcriptomes, we compared gene expression profiles of hybrids and parental species using a quantitative measure of the degree of misexpression [i.e., ∼log_2_(parent-to-hybrid ratio of expression)]. It is important to emphasize that the degree of misexpression of each hybrid genotype was always assessed by comparison with the corresponding parental stocks. For example, the degree of misexpression of attached-X hybrid females was determined based on the comparison to females of the *D. yakuba* and *D. santomea* attached-X parental stocks combined. In this way, we took into account the putative effects of the *Y* chromosome on the transcriptome profiles of attached-X females. To make the female and male transcriptomes comparable, we examined misexpression of autosomal genes only. We found that the hybrid female transcriptome shows a significantly lower degree of misexpression than the hybrid male transcriptome (0.21 *vs*. 0.32; MWT *P* = 1.7×10^-158^ for hybrids from the cross between *D. yakuba* TaïE2 females and *D. santomea* STO.4 males; 0.20 *vs*. 0.34; MWT *P* < 1×10^-308^ for hybrids from the reciprocal cross).

The observation that genes in hybrid males are more prone to misexpression than genes in hybrid females can potentially be attributed to the hemizygosity of the *X* chromosome in males, to faster-male evolution, or a combination of both. To evaluate the contribution of recessive factors on the *X* chromosome to hybrid misexpression, we examined transcript abundance for autosomal genes in attached-X hybrid females (**Figure** 5 for hybrids from the cross between *D. yakuba* TaïE2 females and *D. santomea* STO.4 males). Reminiscent of males, attached-X hybrid females show a significantly greater degree of misexpression than normal hybrid females (0.29 *vs*. 0. 21; MWT *P* = 8.7×10^-71^), with FSGs showing a particularly extreme pattern (**Figure** 6). The increased misexpression in attached-X hybrid females relative to normal hybrid females is also observed in genes with nonsex-biased expression. These results suggest that there is a significant fraction of recessive regulatory incompatibilities involving the *X* chromosome that are masked in normal hybrid females but are fully uncovered in attached-X hybrid females and in hybrid males. The overall degree of misexpression in attached-X hybrid females, however, is not as high as in hybrid males, which suggests that additional factors like faster-male evolution are necessary to fully explain misexpression in hybrid males (0.29 *vs*. 0. 32; MWT *P* = 4.0×10^-23^).

**Figure 5.**
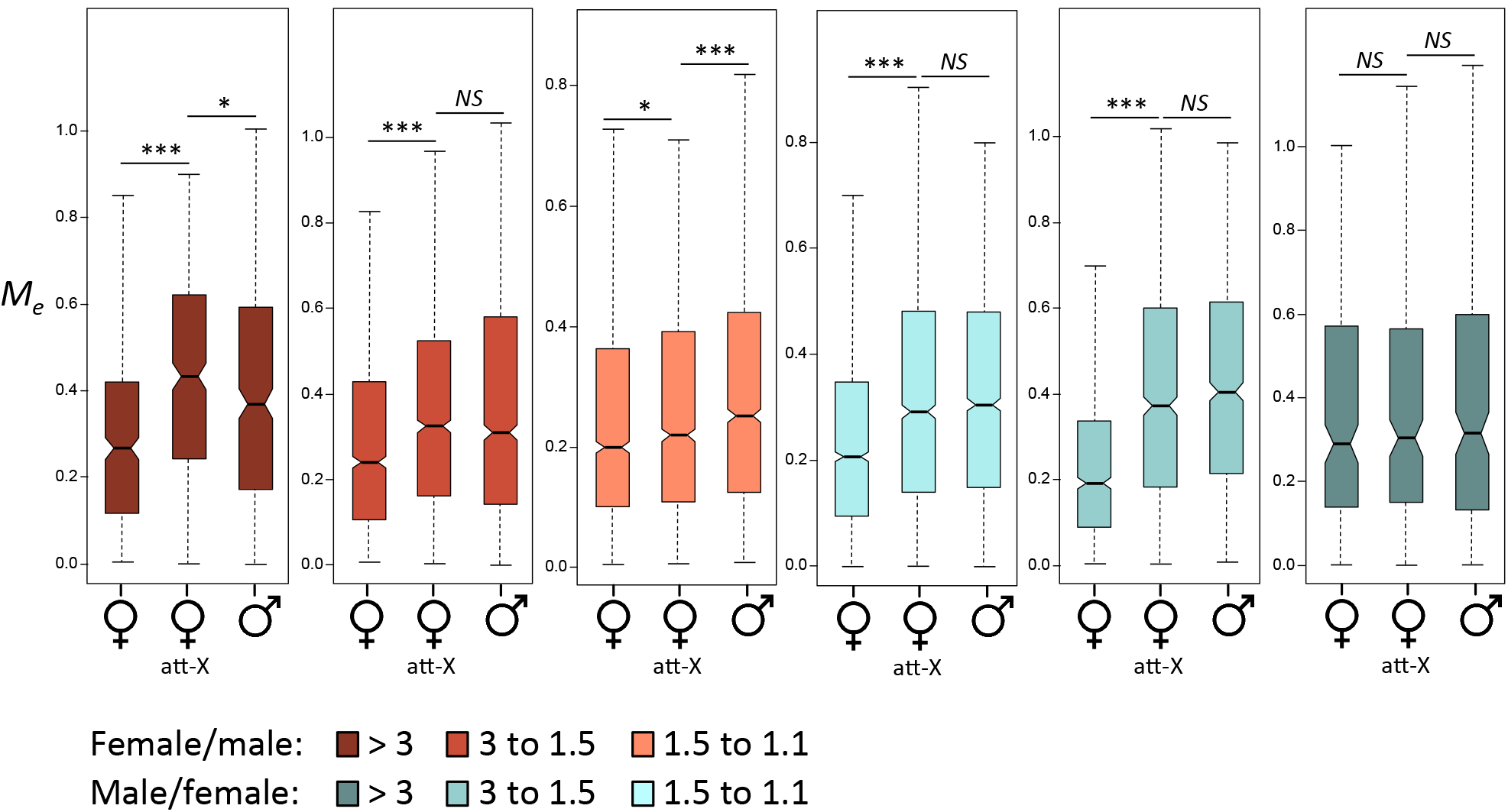
Misexpression (*M_e_*) of autosomal genes with different degrees of sex-biased expression in normal hybrid females, attached-X (att-X) hybrid females and hybrid males. Data based on first generation hybrids from the cross between *D. yakuba* Taï18E2 females and *D. santomea* STO.4 males. Probabilities are based on Mann-Whitney tests where *NS* indicates not significant, \**P* < 0.05, \*\**P* < 0.001, and \*\*\**P* < 1×10^-4^. (See Fig. 3 legend for boxplot explanation.)

**Figure 6.**
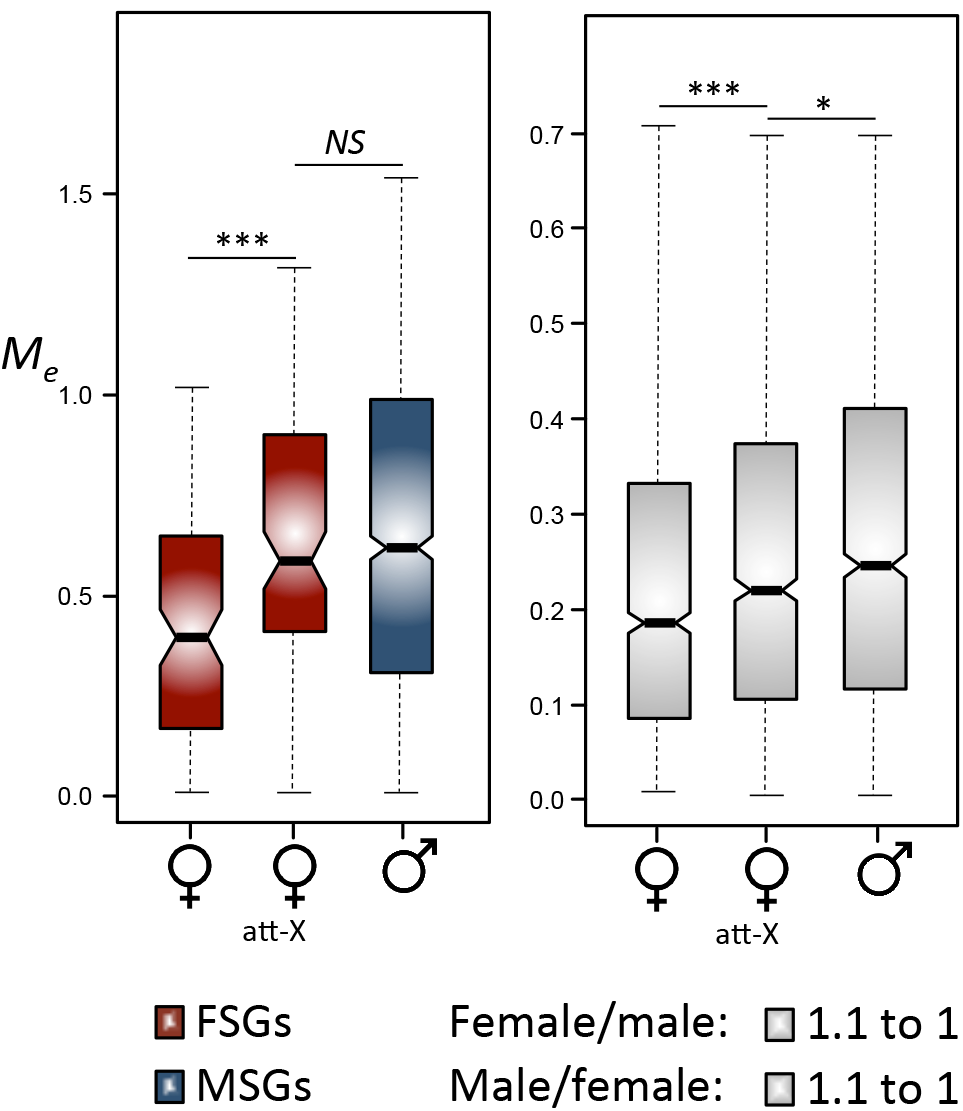
Misexpression (*M_e_*) of autosomal genes with sex-specific and nonsex-biased expression in normal hybrid females, attached-X (att-X) hybrid females and hybrid males. Data based on first generation hybrids from the cross between *D. yakuba* Taï18E2 females and *D. santomea* STO.4 males. FSGs, genes with female-specific expression; MSGs, genes with male-specific expression. Probabilities are based on Mann-Whitney tests where *NS* indicates not significant, \**P* < 0.05, \*\**P* < 0.001, and \*\*\**P* < 1×10^-4^. (See Fig. 3 legend for boxplot explanation.)

To investigate the impact that genetic variation linked to the *X* (and/or *Y*) chromosome may have on gene expression patterns of autosomes, we examined differences in transcript abundance in interspecific hybrids from reciprocal crosses [i.e., ∼log_2_(HFY-to-HFS ratio of expression); **Figure** 1]. Our analysis showed that autosomal genes are more differentially expressed between the two reciprocal hybrid males than between the two attached-X hybrid females (0.20 *vs*. 0.15, MWT *P* = 3.28×10^-80^), even though these females are as ‘unbalanced’ for the *X*-to-autosome ratio as hybrid males. Many of these gene expression differences do not reflect cytoplasmic effects, for the median expression difference between reciprocal hybrid females with normal karyotypes is only 0.10. Remarkably, autosomal MSGs are particularly susceptible to genetic variation linked to sex chromosomes: the median difference in expression between reciprocal hybrid males is almost 8-fold increased relative to the median difference observed for FSGs between reciprocal attached-X hybrid females (1.55 *vs*. 0.20, MWT *P* = 1.35×10^-42^).

## DISCUSSION

Our study provides a comprehensive view of transcriptome evolution in adult females and males of the *D. yakuba*-*D. santomea* system, its molecular mechanisms, and its consequences on patterns of gene expression in interspecific hybrids. Our emphasis lies on the contributions of the *X* chromosome. By taking advantage of attached-X stocks, we have been able to test the dominance theory for hybrid transcriptomes. We have also assessed the effects of linkage to the *X* chromosome on gene expression divergence, which we showed to be influenced by additional factors such as degree of sex-biased expression.

We found that both the female and male transcriptomes show evidence of accelerated gene expression divergence driven by genes with sex-biased expression in the *D. yakuba*-*D. santomea* system. Similar trends have been reported in other systems and have been attributed to relaxed selective constraints due to reduced pleiotropic effects, low levels of expression, and/or a considerable fraction of positive selection, especially in genes with male-biased expression (Meiklejohn *et al.* 2003; Parisi *et al.* 2003; Ranz *et al.* 2003; Zhang *et al.* 2004; Khaitovich *et al.* 2005; Proschel *et al.* 2006; Ellegren and Parsch 2007; Haerty *et al.* 2007; Voolstra *et al.* 2007; Zhang *et al.* 2007; Jiang and Machado 2009; Meisel 2011; Assis *et al.* 2012; Grath and Parsch 2012; Parsch and Ellegren 2013; Harrison *et al.* 2015; Grath and Parsch 2016). Our results also emphasize the importance of studying gene expression divergence in both sexes and reveal the dual nature of transcriptome evolution. While gene expression divergence in genes with sex-biased expression is consistently high in the sex toward which the expression is biased, in the opposite sex transcript abundance appears to be more conserved between species. This low divergence, we propose, reflects strong selective constraints on tissues and pathways shared by both sexes. In contrast, we suggest that the high expression divergence observed for male-biased genes when expressed in males or female-biased genes when expressed in females may be the result of additional sex-specific regulatory components that can potentially evolve by positive selection or under relaxed functional constraints. Our analysis of gene expression divergence on the highly female-biased dot chromosome suggests that weakly deleterious regulatory mutations are major contributors to gene expression divergence of genes with sex-biased expression but only in the sex toward which the expression is biased. Our results also suggest that sexes adopt different expression strategies for their specialized function (Gibilisco *et al.* 2016), possibly through sexually dimorphic *trans*-regulatory environments that often interact with *cis*-regulatory variation differently in females than in males (Coolon *et al.* 2013; Meiklejohn *et al.* 2014).

When we analyzed all the genes expressed in adults as a single class, we found evidence of slower-X evolution of gene expression in both the female the male transcriptomes. The molecular mechanism underlying this increased gene expression divergence in autosomal genes involves contributions from both *cis*- and *trans*-regulatory variation, as proposed for species of the *D. melanogaster* subgroup to explain faster-X evolution of gene expression (Coolon *et al.* 2015). The observations of slower-X evolution can be explained by a significant contribution of weakly deleterious mutations to gene expression divergence, with more effective purifying selection on the *X* chromosome. Alternatively, our findings could be the result of differences in mutation rates between the *X* chromosome and autosomes. In *D. melanogaster* the mutation rate in the male germline is slightly higher than that in the female germline, a difference that can potentially lead to slower-X evolution (Miyata *et al.* 1987; Bachtrog 2008; Keightley *et al.* 2009). There is also a third explanation: in very closely related species ancestral polymorphism represents a non-negligible fraction of divergence and, therefore, slower-X evolution may just reflect the difference in ancestral *N_eX_* and *N_eA_*, with *N_eX_* < *N_eA_*. Consistent with this idea, the genome-wide analysis of putatively neutral sites [bases 8–30 of small introns (< 65 bp);(Llopart 2018)] in the *D. yakuba*-*D. santomea* system, indicates that nucleotide heterozygosity (Watterson 1975) is significantly smaller on the *X* chromosome than on autosomes for a *D. yakuba* population from Nairobi, Kenya, (0.015 *vs*. 0.019; Mann-Whitney test *P* = 0.02) (Rogers *et al.* 2014; Rogers *et al.* 2015). Similarly, a multilocus study of *D. yakuba* and *D. santomea* populations from São Tomé reported that the *N_eX_* /*N_eA_* was close to the theoretical expectation of 0.75 (Llopart *et al.* 2005). As such, a tendency towards slower-X is expected to be the default observation in many closely related species.

Studies of gene expression divergence in several *Drosophila* species show that the chances of observing faster-X (or slower-X) evolution in transcriptome data depend on species, sex, developmental stage, and tissue under study (Khaitovich *et al.* 2005; Brawand *et al.* 2011; Kayserili *et al.* 2012; Llopart 2012; Meisel *et al.* 2012; Meisel and Connallon 2013; Coolon *et al.* 2015; Dean *et al.* 2015). We suggest that this variation may be the result of a combination of biological and experimental factors. Among the biological factors, the *N_eX_* /*N_eA_* has been shown by population genetics models to have a critical impact on whether faster-X evolution is expected (Vicoso and Charlesworth 2009a; Mank *et al.* 2010b). This ratio refers to the long-term, historical *N_eX_* and *N_eA_*, which may not necessarily reflect the estimate based on extant variation. The number of breeding females and males, differences between the two sexes in the variance in reproductive success (i.e., mating system), demographic events, or differences in recombination rates between the *X* chromosome and autosomes (Langley *et al.* 1988; Connallon 2007; Pool and Nielsen 2007; Vicoso and Charlesworth 2009b; Charlesworth 2012; Comeron *et al.* 2012; Avila *et al.* 2015; Comeron 2017) are some of the additional biological factors that can be species specific and considerably affect *N_eX_/N_eA_* (Vicoso and Charlesworth 2009a). It is also important to recognize that factors associated with experimental design, like the specific stocks used in comparisons of closely related species, the subset of genes under study, or statistical cutoffs may also play a critical role in our ability to detect faster-X evolution of gene expression.

Of all the genes with sexually dimorphic expression analyzed in this study, X-linked MSGs are the most distinct class. They show transcriptional faster-X and their rampant gene expression divergence is even greater than that of the rapidly evolving female-biased genes on the dot chromosome. Based on neutral polymorphism surveys, there is a 5-10 fold reduction in effective population size and, therefore, in the scaled-population selection coefficient on the dot chromosome relative to the *X* and autosomes (Berry *et al.* 1991; Jensen *et al.* 2002; Wang *et al.* 2002; Wang *et al.* 2004; Betancourt *et al.* 2009; Arguello *et al.* 2010). As a result, we suggest that it is unlikely that the faster-X gene expression evolution of MSGs is due to relaxed selection against deleterious mutations on the *X* chromosome. The overall finding of slower-X evolution for male transcriptomes also militates against this possibility. Instead, we propose that the faster-X observed for these genes in the *D. yakuba*-*D. santomea* system is the result of a higher incidence of beneficial regulatory mutations, perhaps associated with sexual selection and/or sexual conflict (Parsch and Ellegren 2013).

Our examination of gene expression in attached-X hybrid females shows that misexpression increases when the *X* chromosome is made homozygous. It is important to emphasize that this is not necessarily an inevitable result because, by carrying two different *X* chromosomes, normal hybrid females are expected to suffer twice as many possible hybrid incompatibilities involving the *X* chromosome as hybrid males (Orr 1993; Orr and Turelli 1996). It is also important to point out that none of the misexpression observed in attached-X hybrid females in the *D. yakuba*-*D. santomea* system is associated with sterility because these females are mostly fertile. Our results, however, suggest that there is a substantial fraction of recessive mutations that have the potential to generate regulatory incompatibilities involving the *X* chromosome. They open the possibility that the hemizygosity of the *X* chromosome in males may significantly contribute to misexpression and ultimately lead to hybrid male sterility. Thus, regulatory incompatibilities in hybrid transcriptomes appear to conform to the dominance theory in the *D. yakuba*-*D. santomea* system (Muller 1940; Muller 1942).

Both dominance theory and faster-X evolution have been proposed as forces that can explain the disproportionally large contribution of the *X* chromosome to hybrid dysfunction, particularly hybrid male sterility (i.e., the large X-effect; COYNE AND ORR 1989) (Turelli and Orr 1995; Orr and Turelli 1996; Turelli and Orr 2000; Masly and Presgraves 2007; Presgraves 2008; Moyle *et al.* 2010). As the rate of evolution of X-linked loci exceeds that of autosomal loci, the large X-effect becomes more likely (Coyne and Orr 1989; Naveira 2003). Similarly, if alleles decreasing fitness in hybrids tend to act recessively (i.e., dominance theory), it is easier to explain the large X-effect, even when the rates of evolution for the *X* chromosome and autosomes are not different (Turelli and Orr 1995; Orr and Turelli 1996; Turelli and Orr 2000). While either force alone can generate the large X-effect, in species like *D. yakuba* and *D. santomea* the two forces pull together in the same direction for genes with a high degree of male-biased expression to easily yield the observed large X-effect. These arguments echo those put forward by Coyne and Orr (2004) to explain the relative contributions of dominance theory and faster-male evolution to Haldane’s rule.

In the context of gene expression, our results suggest that recessive X-linked mutations and faster-X evolution of MSGs can potentially play a major role in hybrid male misexpression and perhaps the large X-effect. Our results indicate that genetic variation linked to sex chromosomes substantially impacts the expression levels of *autosomal* MSGs. Although we have not formally evaluated regulatory interactions, these observations open the possibility that regulatory incompatibilities between X-linked and autosomal factors may be facilitating the large X-effect on hybrid male sterility.

## REFERENCES

Afgan, E., D. Baker, M. van den Beek, D. Blankenberg, D. Bouvier et al., 2016 The Galaxy platform for accessible, reproducible and collaborative biomedical analyses: 2016 update. Nucleic Acids Res 44: W3–W10.

Arguello, J. R., Y. Zhang, T. Kado, C. Fan, R. Zhao et al., 2010 Recombination yet inefficient selection along the *Drosophila melanogaster* subgroup’s fourth chromosome. Mol Biol Evol 27: 848–861.

Artieri, C. G., W. Haerty and R. S. Singh, 2007 Association between levels of coding sequence divergence and gene misregulation in *Drosophila* male hybrids. J Mol Evol 65: 697–704.

Assis, R., Q. Zhou and D. Bachtrog, 2012 Sex-biased transcriptome evolution in Drosophila. Genome Biol Evol 4: 1189–1200.

Avila, V., J. L. Campos and B. Charlesworth, 2015 The effects of sex-biased gene expression and X-linkage on rates of adaptive protein sequence evolution in *Drosophila*. Biol Lett 11: 20150117.

Avila, V., S. Marion de Proce, J. L. Campos, H. Borthwick, B. Charlesworth et al., 2014 Faster-X effects in two *Drosophila* lineages. Genome Biol Evol 6: 2968–2982.

Bachtrog, D., 2008 Evidence for male-driven evolution in Drosophila. Mol Biol Evol 25: 617–619.

Baines, J. F., and B. Harr, 2007 Reduced X-linked diversity in derived populations of house mice. Genetics 175: 1911–1921.

Baines, J. F., S. A. Sawyer, D. L. Hartl and J. Parsch, 2008 Effects of X-linkage and sex-biased gene expression on the rate of adaptive protein evolution in *Drosophila*. Mol Biol Evol 25: 1639–1650.

Barbash, D. A., and J. G. Lorigan, 2007 Lethality in *Drosophila melanogaster/Drosophila simulans* species hybrids is not associated with substantial transcriptional misregulation. J Exp Zoolog B Mol Dev Evol 308: 74–84.

Begun, D. J., A. K. Holloway, K. Stevens, L. W. Hillier, Y. P. Poh et al., 2007 Population genomics: whole-genome analysis of polymorphism and divergence in *Drosophila simulans*. PLoS Biol. 5: e310.

Benjamini, Y., and Y. Hochberg, 1995 Controlling the false discovery rate: a practical and powerful approach to multiple testing. J. R. Stat. Soc. B 57: 289–300.

Berry, A. J., J. W. Ajioka and M. Kreitman, 1991 Lack of polymorphism on the *Drosophila* fourth chromosome resulting from selection. Genetics 129: 1111–1117.

Betancourt, A. J., and D. C. Presgraves, 2002 Linkage limits the power of natural selection in *Drosophila*. Proc. Natl. Acad. Sci. USA 99: 13616–13620.

Betancourt, A. J., D. C. Presgraves and W. J. Swanson, 2002 A Test for Faster X Evolution in *Drosophila*. Mol. Biol. Evol. 19: 1816–1819.

Betancourt, A. J., J. J. Welch and B. Charlesworth, 2009 Reduced effectiveness of selection caused by a lack of recombination. Curr Biol 19: 655–660.

Brawand, D., M. Soumillon, A. Necsulea, P. Julien, G. Csardi et al., 2011 The evolution of gene expression levels in mammalian organs. Nature 478: 343–348.

Catron, D. J., and M. A. Noor, 2008 Gene expression disruptions of organism versus organ in *Drosophila* species hybrids. PLoS ONE 3: e3009.

Charlesworth, B., 2012 The role of background selection in shaping patterns of molecular evolution and variation: evidence from variability on the *Drosophila* X chromosome. Genetics 191: 233–246.

Charlesworth, B., J. A. Coyne and N. Barton, 1987 The relative rates of evolution of sex chromosomes and autosomes. Am. Nat. 130: 113–146.

Civetta, A., 2016 Misregulation of Gene Expression and Sterility in Interspecies Hybrids: Causal Links and Alternative Hypotheses. J Mol Evol 82: 176–182.

Clark, A. G., M. B. Eisen, D. R. Smith, C. M. Bergman, B. Oliver et al., 2007 Evolution of genes and genomes on the *Drosophila* phylogeny. Nature 450: 203–218.

Comeron, J. M., 2017 Background selection as null hypothesis in population genomics: Insights and challenges from *Drosophila* studies. Philosophical Transactions of the Royal Society B in press.

Comeron, J. M., and M. Kreitman, 2002 Population, evolutionary and genomic consequences of interference selection. Genetics 161: 389–410.

Comeron, J. M., R. Ratnappan and S. Bailin, 2012 The many landscapes of recombination in *Drosophila melanogaster*. PLoS Genetics 8: e1002905.

Comeron, J. M., A. Williford and R. M. Kliman, 2008 The Hill-Robertson effect: evolutionary consequences of weak selection and linkage in finite populations. Heredity (Edinb) 100: 19–31.

Connallon, T., 2007 Adaptive protein evolution of X-linked and autosomal genes in *Drosophila* : implications for faster-X hypotheses. Mol Biol Evol 24: 2566–2572.

Coolon, J. D., C. J. McManus, K. R. Stevenson, B. R. Graveley and P. J. Wittkopp, 2014 Tempo and mode of regulatory evolution in *Drosophila*. Genome Res 24: 797–808.

Coolon, J. D., K. R. Stevenson, C. J. McManus, B. Yang, B. R. Graveley et al., 2015 Molecular Mechanisms and Evolutionary Processes Contributing to Accelerated Divergence of Gene Expression on the *Drosophila* X Chromosome. Mol Biol Evol 32: 2605–2615.

Coolon, J. D., W. Webb and P. J. Wittkopp, 2013 Sex-specific effects of cis-regulatory variants in *Drosophila melanogaster*. Genetics 195: 1419–1422.

Counterman, B. A., D. Ortiz-Barrientos and M. A. Noor, 2004 Using comparative genomic data to test for fast-X evolution. Evolution 58: 656–660.

Coyne, J. A., 1992 Genetics and speciation. Nature 355: 511–515.

Coyne, J. A., S. Elwyn, S. Y. Kim and A. Llopart, 2004 Genetic studies of two sister species in the *Drosophila melanogaster* subgroup, *D. yakuba* and *D. santomea*. Genetical Research 84: 11–26.

Coyne, J. A., and H. A. Orr, 1989 Two rules of speciation, pp. 180–207 in Speciation and its Consequences, edited by D. Otte and J. Endler. Sinauer Associates, Sunderland, MA.

Coyne, J. A., and H. A. Orr, 2004 Speciation. Sinauer Associates, Sunderland, MA.

Coyne, J. A., J. Rux and J. R. David, 1991 Genetics of morphological differences and hybrid sterility between *Drosophila sechellia* and its relatives. Genet Res 57: 113–122.

Dean, R., P. W. Harrison, A. E. Wright, F. Zimmer and J. E. Mank, 2015 Positive Selection Underlies Faster-Z Evolution of Gene Expression in Birds. Mol Biol Evol 32: 2646–2656.

Dobzhansky, T., 1937 Genetics and the origin of species. Columbia University Press, New York.

Ellegren, H., 2009 Genomic evidence for a large-Z effect. Proc Biol Sci 276: 361–366.

Ellegren, H., and J. Parsch, 2007 The evolution of sex-biased genes and sex-biased gene expression. Nat Rev Genet 8: 689–698.

Garrigan, D., S. B. Kingan, A. J. Geneva, J. P. Vedanayagam and D. C. Presgraves, 2014 Genome Diversity and Divergence in *Drosophila mauritiana* : Multiple Signatures of Faster X Evolution. Genome Biol Evol 6: 2444–2458.

Gibilisco, L., Q. Zhou, S. Mahajan and D. Bachtrog, 2016 Alternative Splicing within and between Drosophila Species, Sexes, Tissues, and Developmental Stages. PLoS Genet 12: e1006464.

Gomes, S., and A. Civetta, 2015 Hybrid male sterility and genome-wide misexpression of male reproductive proteases. Sci Rep 5: 11976.

Gramates, L. S., S. J. Marygold, G. D. Santos, J. M. Urbano, G. Antonazzo et al., 2017 FlyBase at 25: looking to the future. Nucleic Acids Res 45: D663–D671.

Grath, S., and J. Parsch, 2012 Rate of amino acid substitution is influenced by the degree and conservation of male-biased transcription over 50 million years of *Drosophila* evolution. Genome Biol Evol 4: 346–359.

Grath, S., and J. Parsch, 2016 Sex-Biased Gene Expression. Annu Rev Genet 50: 29–44.

Guerrero, R. F., C. D. Muir, S. Josway and L. C. Moyle, 2017 Pervasive antagonistic interactions among hybrid incompatibility loci. PLoS Genet 13: e1006817.

Haddrill, P. R., D. L. Halligan, D. Tomaras and B. Charlesworth, 2007 Reduced efficacy of selection in regions of the Drosophila genome that lack crossing over. Genome Biol 8: R18.

Haerty, W., S. Jagadeeshan, R. J. Kulathinal, A. Wong, K. Ravi Ram et al., 2007 Evolution in the fast lane: rapidly evolving sex-related genes in *Drosophila*. Genetics 177: 1321–1335.

Haerty, W., and R. S. Singh, 2006 Gene Regulation Divergence is a Major Contributor to the Evolution of Dobzhansky-Muller Incompatibilities between Species of *Drosophila*. Mol Biol Evol 23: 1707–1714.

Harrison, P. W., A. E. Wright, F. Zimmer, R. Dean, S. H. Montgomery et al., 2015 Sexual selection drives evolution and rapid turnover of male gene expression. Proc Natl Acad Sci U S A 112: 4393–4398.

Hill, W. G., and A. Robertson, 1966 The effect of linkage on limits to artificial selection. Genet. Res. 8: 269–294.

Hu, T. T., M. B. Eisen, K. R. Thornton and P. Andolfatto, 2013 A second-generation assembly of the *Drosophila simulans* genome provides new insights into patterns of lineage-specific divergence. Genome Res 23: 89–98.

Jensen, M. A., B. Charlesworth and M. Kreitman, 2002 Patterns of genetic variation at a chromosome 4 locus of Drosophila melanogaster and D. simulans. Genetics 160: 493–507.

Jiang, Z. F., and C. A. Machado, 2009 Evolution of sex-dependent gene expression in three recently diverged species of *Drosophila*. Genetics 183: 1175–1185.

Kayserili, M. A., D. T. Gerrard, P. Tomancak and A. T. Kalinka, 2012 An excess of gene expression divergence on the X chromosome in *Drosophila* embryos: implications for the faster-X hypothesis. PLoS Genet 8: e1003200.

Keightley, P. D., U. Trivedi, M. Thomson, F. Oliver, S. Kumar et al., 2009 Analysis of the genome sequences of three Drosophila melanogaster spontaneous mutation accumulation lines. Genome Res 19: 1195–1201.

Khaitovich, P., I. Hellmann, W. Enard, K. Nowick, M. Leinweber et al., 2005 Parallel patterns of evolution in the genomes and transcriptomes of humans and chimpanzees. Science 309: 1850–1854.

Kousathanas, A., D. L. Halligan and P. D. Keightley, 2014 Faster-X adaptive protein evolution in house mice. Genetics 196: 1131–1143.

Lachaise, D., M. Harry, M. Solignac, F. Lemeunier, V. Bénassi et al., 2000 Evoutionary novelties in islands: *Drosophila santomea* a new *melanogaster* sister species from São Tomé. Proc. Roy. Soc. Lond. B 267: 1487–1495.

Landry, C. R., P. J. Wittkopp, C. H. Taubes, J. M. Ranz, A. G. Clark et al., 2005 Compensatory cis-trans evolution and the dysregulation of gene expression in interspecific hybrids of *Drosophila*. Genetics 171: 1813–1822.

Langley, C. H., E. Montgomery, R. Hudson, N. Kaplan and B. Charlesworth, 1988 On the role of unequal exchange in the containment of transposable element copy number. Genet. Res. 52: 223–235.

Langmead, B., and S. L. Salzberg, 2012 Fast gapped-read alignment with Bowtie 2. Nat Methods 9: 357–359.

Larson, E. L., D. Vanderpool, S. Keeble, M. Zhou, B. A. Sarver et al., 2016 Contrasting Levels of Molecular Evolution on the Mouse X Chromosome. Genetics 203: 1841–1857.

Li, B., and C. N. Dewey, 2011 RSEM: accurate transcript quantification from RNA-Seq data with or without a reference genome. BMC Bioinformatics 12: 323.

Li, H., B. Handsaker, A. Wysoker, T. Fennell, J. Ruan et al., 2009 The Sequence Alignment/Map format and SAMtools. Bioinformatics 25: 2078–2079.

Llopart, A., 2012 The rapid evolution of X-linked male-biased gene expression and the large-X effect in *Drosophila yakuba, D*. santomea and their hybrids. Mol. Biol. Evol. 29: 3873–3886.

Llopart, A., 2015 Parallel faster-X evolution of gene expression and protein sequences in *Drosophila* : beyond differences in expression properties and protein interactions. PLoS ONE 10: e0116829.

Llopart, A., 2018 Faster-X evolution of gene expression is driven by recessive adaptive cis-regulatory variation in Drosophila. Mol Ecol. (in press)

Llopart, A., D. Lachaise and J. A. Coyne, 2005 Multilocus analysis of introgression between two sympatric sister species of *Drosophila* : *Drosophila yakuba* and *D. santomea*. Genetics 171: 197–210.

Love, M. I., W. Huber and S. Anders, 2014 Moderated estimation of fold change and dispersion for RNA-seq data with DESeq2. Genome Biol 15: 550.

Lu, J., and C. I. Wu, 2005 Weak selection revealed by the whole-genome comparison of the X chromosome and autosomes of human and chimpanzee. Proc Natl Acad Sci U S A 102: 4063–4067.

Maheshwari, S., and D. A. Barbash, 2012 Cis-by-Trans regulatory divergence causes the asymmetric lethal effects of an ancestral hybrid incompatibility gene. PLoS Genet 8: e1002597.

Mank, J. E., E. Axelsson and H. Ellegren, 2007 Fast-X on the Z: rapid evolution of sex-linked genes in birds. Genome Res 17: 618–624.

Mank, J. E., K. Nam and H. Ellegren, 2010a Faster-Z evolution is predominantly due to genetic drift. Mol Biol Evol 27: 661–670.

Mank, J. E., B. Vicoso, S. Berlin and B. Charlesworth, 2010b Effective population size and the Faster-X effect: empirical results and their interpretation. Evolution 64: 663–674.

Masly, J. P., and D. C. Presgraves, 2007 High-resolution genome-wide dissection of the two rules of speciation in *Drosophila*. PLoS Biol 5: e243.

Matute, D. R., I. A. Butler, D. A. Turissini and J. A. Coyne, 2010 A test of the snowball theory for the rate of evolution of hybrid incompatibilities. Science 329: 1518–1521.

McManus, C. J., J. D. Coolon, M. O. Duff, J. Eipper-Mains, B. R. Graveley et al., 2010 Regulatory divergence in Drosophila revealed by mRNA-seq. Genome Res 20: 816–825.

Meiklejohn, C. D., J. D. Coolon, D. L. Hartl and P. J. Wittkopp, 2014 The roles of cis-and trans-regulation in the evolution of regulatory incompatibilities and sexually dimorphic gene expression. Genome Res 24: 84–95.

Meiklejohn, C. D., J. Parsch, J. M. Ranz and D. L. Hartl, 2003 Rapid evolution of male-biased gene expression in *Drosophila*. Proc Natl Acad Sci U S A 100: 9894–9899.

Meisel, R. P., 2011 Towards a more nuanced understanding of the relationship between sex-biased gene expression and rates of protein-coding sequence evolution. Mol Biol Evol 28: 1893–1900.

Meisel, R. P., and T. Connallon, 2013 The faster-X effect: integrating theory and data. Trends Genet 29: 537–544.

Meisel, R. P., J. H. Malone and A. G. Clark, 2012 Faster-X evolution of gene expression in *Drosophila*. PLoS Genet. 8: e1003013.

Michalak, P., and M. A. Noor, 2003 Genome-wide patterns of expression in *Drosophila* pure species and hybrid males. Mol Biol Evol 20: 1070–1076.

Michalak, P., and M. A. Noor, 2004 Association of misexpression with sterility in hybrids of *Drosophila simulans* and *D. mauritiana*. J Mol Evol 59: 277–282.

Miyata, T., H. Hayashida, K. Kuma, K. Mitsuyasu and T. Yasunaga, 1987 Male-driven molecular evolution: a model and nucleotide sequence analysis. Cold Spring Harb Symp Quant Biol 52: 863–867.

Moehring, A. J., K. C. Teeter and M. A. Noor, 2007 Genome-wide patterns of expression in *Drosophila* pure species and hybrid males. II. Examination of multiple-species hybridizations, platforms, and life cycle stages. Mol Biol Evol 24: 137–145.

Moyle, L. C., C. D. Muir, M. V. Han and M. W. Hahn, 2010 The contribution of gene movement to the “two rules of speciation”. Evolution 64: 1541–1557.

Moyle, L. C., and T. Nakazato, 2010 Hybrid incompatibility “snowballs” between Solanum species. Science 329: 1521–1523.

Muller, H. J., 1940 Bearing of the *Drosophila* work on systematics, pp. 185–268 in The New Systematics., edited by J. S. Huxley. Clarendon Press, Oxford.

Muller, H. J., 1942 Isolating mechanisms, evolution, and temperature. Biol. Symp. 6: 71–125.

Musters, H., M. A. Huntley and R. S. Singh, 2006 A genomic comparison of faster-sex, faster-X, and faster-male evolution between *Drosophila melanogaster* and *Drosophila pseudoobscura*. J Mol Evol 62: 693–700.

Naveira, H. F., 2003 On the relative roles of faster-X evolution and dominance in the establishment of intrinsic postzygotic isolating barriers. Genetica 118: 41–50.

Nielsen, R., C. Bustamante, A. G. Clark, S. Glanowski, T. B. Sackton et al., 2005 A scan for positively selected genes in the genomes of humans and chimpanzees. PLoS Biol 3: e170.

Noor, M. A., 2005 Patterns of evolution of genes disrupted in expression in *Drosophila* species hybrids. Genet Res 85: 119–125.

Orr, H. A., 1987 Genetics of male and female sterility in hybrids of *Drosophila pseudoobscura* and *D. persimilis*. Genetics 116: 555–563.

Orr, H. A., 1993 A mathematical model of Haldane’s rule. Evolution 47: 1606–1611.

Orr, H. A., 1995 The population genetics of speciation: the evolution of hybrid incompatibilities. Genetics 139: 1805–1813.

Orr, H. A., and A. J. Betancourt, 2001 Haldane’s sieve and adaptation from the standing genetic variation. Genetics 157: 875–884.

Orr, H. A., and M. A. Turelli, 1996 Dominance and Haldane’s rule. Genetics 143: 613–616.

Ortiz-Barrientos, D., B. A. Counterman and M. A. Noor, 2007 Gene expression divergence and the origin of hybrid dysfunctions. Genetica 129: 71–81.

Parisi, M., R. Nuttall, D. Naiman, G. Bouffard, J. Malley et al., 2003 Paucity of genes on the *Drosophila* X chromosome showing male-biased expression. Science 299: 697–700.

Parsch, J., and H. Ellegren, 2013 The evolutionary causes and consequences of sex-biased gene expression. Nat Rev Genet 14: 83–87.

Pool, J. E., and R. Nielsen, 2007 Population size changes reshape genomic patterns of diversity. Evolution 61: 3001–3006.

Presgraves, D. C., 2008 Sex chromosomes and speciation in *Drosophila*. Trends Genet 24: 336–343.

Proschel, M., Z. Zhang and J. Parsch, 2006 Widespread adaptive evolution of *Drosophila* genes with sex-biased expression. Genetics 174: 893–900.

Ranz, J. M., C. I. Castillo-Davis, C. D. Meiklejohn and D. L. Hartl, 2003 Sex-Dependent Gene Expression and Evolution of the *Drosophila* Transcriptome. Science 300: 1742–1745.

Ranz, J. M., K. Namgyal, G. Gibson and D. L. Hartl, 2004 Anomalies in the expression profile of interspecific hybrids of *Drosophila melanogaster* and *Drosophila simulans*. Genome Res 14: 373–379.

Reiland, J., and M. A. Noor, 2002 Little qualitative RNA misexpression in sterile male F1 hybrids of *Drosophila pseudoobscura* and *D. persimilis*. BMC Evol Biol 2: 16.

Rogers, R. L., J. M. Cridland, L. Shao, T. T. Hu, P. Andolfatto et al., 2014 Landscape of standing variation for tandem duplications in Drosophila yakuba and Drosophila simulans. Mol Biol Evol 31: 1750–1766.

Rogers, R. L., J. M. Cridland, L. Shao, T. T. Hu, P. Andolfatto et al., 2015 Tandem Duplications and the Limits of Natural Selection in *Drosophila yakuba* and *Drosophila simulans*. PLoS One 10: e0132184.

Sackton, T. B., R. B. Corbett-Detig, J. Nagaraju, L. Vaishna, K. P. Arunkumar et al., 2014 Positive Selection Drives Faster-Z Evolution in Silkmoths. Evolution.

Satyaki, P. R., T. N. Cuykendall, K. H. Wei, N. J. Brideau, H. Kwak et al., 2014 The Hmr and Lhr hybrid incompatibility genes suppress a broad range of heterochromatic repeats. PLoS Genet 10: e1004240.

Singh, N. D., A. M. Larracuente and A. G. Clark, 2008 Contrasting the efficacy of selection on the X and autosomes in *Drosophila*. Mol Biol Evol 25: 454–467.

Sundararajan, V., and A. Civetta, 2011 Male sex interspecies divergence and down regulation of expression of spermatogenesis genes in *Drosophila* sterile hybrids. J Mol Evol 72: 80–89.

Thornton, K., and M. Long, 2002 Rapid divergence of gene duplicates on the *Drosophila melanogaster* X chromosome. Mol Biol Evol 19: 918–925.

Thornton, K., and M. Long, 2005 Excess of amino acid substitutions relative to polymorphism between X-linked duplications in *Drosophila melanogaster*. Mol Biol Evol 22: 273–284.

Torgerson, D. G., and R. S. Singh, 2003 Sex-linked mammalian sperm proteins evolve faster than autosomal ones. Mol Biol Evol 20: 1705–1709.

Torgerson, D. G., and R. S. Singh, 2006 Enhanced adaptive evolution of sperm-expressed genes on the mammalian X chromosome. Heredity (Edinb) 96: 39–44.

Turelli, M., and H. A. Orr, 1995 The dominance theory of Haldane’s rule. Genetics 140: 389–402.

Turelli, M., and H. A. Orr, 2000 Dominance, epistasis and the genetics of postzygotic isolation. Genetics 154: 1663–1679.

Veeramah, K. R., R. N. Gutenkunst, A. E. Woerner, J. C. Watkins and M. F. Hammer, 2014 Evidence for increased levels of positive and negative selection on the X chromosome versus autosomes in humans. Mol Biol Evol.

Vicoso, B., and D. Bachtrog, 2013 Reversal of an ancient sex chromosome to an autosome in Drosophila. Nature 499: 332–335.

Vicoso, B., and D. Bachtrog, 2015 Numerous transitions of sex chromosomes in Diptera. PLoS Biol 13: e1002078.

Vicoso, B., and B. Charlesworth, 2006 Evolution on the X chromosome: unusual patterns and processes. Nat Rev Genet 7: 645–653.

Vicoso, B., and B. Charlesworth, 2009a Effective population size and the faster-X effect: an extended model. Evolution 63: 2413–2426.

Vicoso, B., and B. Charlesworth, 2009b Recombination rates may affect the ratio of X to autosomal noncoding polymorphism in African populations of *Drosophila melanogaster*. Genetics 181: 1699–1701.

Vicoso, B., J. J. Emerson, Y. Zektser, S. Mahajan and D. Bachtrog, 2013 Comparative sex chromosome genomics in snakes: differentiation, evolutionary strata, and lack of global dosage compensation. PLoS Biol 11: e1001643.

Voolstra, C., D. Tautz, P. Farbrother, L. Eichinger and B. Harr, 2007 Contrasting evolution of expression differences in the testis between species and subspecies of the house mouse. Genome Res 17: 42–49.

Wang, W., K. Thornton, A. Berry and M. Long, 2002 Nucleotide variation along the *Drosophila melanogaste*r fourth chromosome. Science 295: 134–137.

Wang, W., K. Thornton, J. J. Emerson and M. Long, 2004 Nucleotide variation and recombination along the fourth chromosome in Drosophila simulans. Genetics 166: 1783–1794.

Watterson, G. A., 1975 On the number of segregating sites in genetical models without recombination. Theor. Popul. Biol. 7: 256–276.

Wei, K. H., A. G. Clark and D. A. Barbash, 2014 Limited gene misregulation is exacerbated by allele-specific upregulation in lethal hybrids between *Drosophila melanogaster* and *Drosophila simulans*. Mol Biol Evol 31: 1767–1778.

Wittkopp, P. J., B. K. Haerum and A. G. Clark, 2004 Evolutionary changes in *cis* and *trans* gene regulation. Nature 430: 85–88.

Wu, C.-I., and A. W. Davis, 1993 Evolution of postmating reproductive isolation: the composite nature of Haldane’s rule and its genetic bases. Am. Nat. 142: 187–212.

Wu, C. I., N. Johnson and M. F. Palopoli, 1996 Haldane’s rule and its legacy: why are there so many sterile males. Trends. Ecol. Evol. 11: 281–284.

Zhang, Y., D. Sturgill, M. Parisi, S. Kumar and B. Oliver, 2007 Constraint and turnover in sex-biased gene expression in the genus *Drosophila*. Nature 450: 233–237.

Zhang, Z., T. M. Hambuch and J. Parsch, 2004 Molecular evolution of sex-biased genes in *Drosophila*. Mol Biol Evol 21: 2130–2139.

